# Achieving High-Resolution Whole-Brain Slab ^1^H-MRSI with Compressed-Sensing and Low-Rank Reconstruction at 7 Tesla

**DOI:** 10.1101/2020.05.13.092668

**Authors:** Antoine Klauser, Bernhard Strasser, Bijaya Thapa, Francois Lazeyras, Ovidiu Andronesi

## Abstract

Low sensitivity MR techniques such as magnetic resonance spectroscopic imaging (MRSI) greatly benefit from the gain in signal-to-noise (SNR) provided by ultra-high field MR. High-resolution and whole-brain slab MRSI remains however very challenging due to lengthy acquisition, low signal, lipid contamination and field inhomogeneity. In this study, we propose an acquisition-reconstruction scheme that combines a ^1^H-FID-MRSI sequence with compressed sensing acceleration and low-rank modeling with total-generalized-variation constraint to achieve metabolite imaging in two and three dimensions at 7 Tesla. The resulting images and volumes reveal highly detailed distributions that are specific to each metabolite and follow the underlying brain anatomy. The MRSI method was validated in a high-resolution phantom containing fine metabolite structures, and in 3 healthy volunteers. This new application of compressed sensing acceleration paves the way for high-resolution MRSI in clinical setting with acquisition times of 5 min for 2D MRSI at 2.5 mm and of 20 min for 3D MRSI at 3.3 mm isotropic.

## I. INTRODUCTION

Magnetic resonance spectroscopy (MRS) [1] has been one of the main motivations driving MR towards ultra-high field (≥ 7T). With the recent advent of FDA approval and CE certification of 7T MR systems for clinical use, there is high interest in developing robust fast high-resolution MRS imaging (MRSI) methods to map the neurochemistry of human brain with greater anatomical details. MRS at 7T and higher field, benefits from increased sensitivity and spectral dispersion [2, 3], which are the two intrinsic factors limiting the chemical information that can be extracted at the current clinical (*≤* 3T) fields. However, capitalizing the benefits of ultra-high field have proven more challenging for human MRS imaging (MRSI) compared to single voxel spectroscopy (SVS) [4]. Large *B*_0_ and *B*_1_ inhomogeneity over the human brain [5, 6] results in non-uniform image quality across an extended field of view. Acquisition of the 4D (k,t) space by traditional clinical MRSI sequences [7] is slow due to long repetition time (TR) needed to sample adequately the time domain, and due to notably high specific-absorption rate (SAR) at ultra-high field which further limits the minimum TR. These problems prohibit the acquisition of spatial high-resolution MRSI to benefit from the higher signal-to-noise (SNR) at ultra-high field. Significant progress has been done in the recent years in accelerating high-resolution MRSI at ultra-high field [8] by employing sequences with low SAR, short TR and short echo time (TE). In particular, the pulse acquire (FID) excitation [9, 10] has grown in popularity at ultra-high field due to the robustness of low flip angle excitation to *B*_1_ transmit inhomogeneity, whereas reduced SAR allows for short TR, and the very short TE reduces signal loss due to 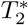 relaxation. The elimination of fat suppression pulses further reduces SAR and TR at ultra-high field but requires advanced reconstruction methods to decontaminate metabolic signal from skull lipid leakage [11–14]. The problem of MRSI acquisition time at ultra-high field was addressed with the use of spectro-spatial encoding using echo-planar [15], concentric circles [16], rosettes [17] and spiral [18] trajectories, but also with parallel imaging methods [19, 20], and more recently with a combination of non-Cartesian trajectories and parallel imaging [21].

Compressed sensing (CS) is a technique that allows for faithful recovery of an undersampled signal [22]. Its application to MRI enable acceleration of the acquisition by random k-space undersampling [23, 24]. Although now being widely used in clinics [25, 26] for standard MRI, CS has been scarcely explored for ^1^H-MRSI in humans, mostly at 3T [27–30] and in one study for single slice 2D MRSI at 9.4T [31]. However, 2D MRSI is typically performed with thick slices (8-10 mm) which lead to partial volume averaging, while isotropic high-resolution 3D coverage is preferable to match better the fine details of gray/white matter structure. Initial CS demonstrations of ^1^H-MRSI at 3T have been shown with volume localization [27–30] which excludes the scalp to avoid lipid artifacts, but also misses most of the lateral cortex. Compressed sensing has been successfully used to speed-up acquisition of X-nuclei MRSI with full field of view (FOV) coverage, such as ^13^C [32] and ^31^P [33], where spectra are more sparse, the large nuisance signals of fat and water are absent, and the effect of *B*_0_ inhomogeneity is reduced. On the other hand for whole brain CS ^1^H-MRSI at ultra-high field, proper separation between metabolites and dominant fat and water signals is of paramount importance [34, 35]. Compared to parallel imaging, the random undersampling in CS has the advantage that it does not produce structured aliasing artefacts, is not affected by the g-factor penalty, does not requires a calibration scan [23], but instead involves a more lengthy reconstruction which is acceptable when a real-time or fast answer is not needed such as often is the case for MRSI, although clinically acceptable reconstruction times can be obtained with optimized algorithms [36].

In this research we investigate the potential of CS combined with a low-rank (LR) constraint and sensitivity encoding (SENSE) in achieving fast high-resolution 2D and isotropic 3D ^1^H-MRSI at 7T for a whole slab of human brain. The ability to accelerate single slice 2D MRSI using CS-SENSE reconstruction was demonstrated at 9.4T by Nassirpour *et al*. for an FID phase encoded sequence [31] and at 3T by Otazo *et al*. for a spin-echo PEPSI sequence [27] or by Chatnuntawech *et al*. for PRESS with random spiral sequence [29]. On the other hand, spectral-spatial separability exploited by low-rank modelling has been shown to be very effective to improve SNR and remove nuisance signal, enabling whole brain high-resolution ^1^H-MRSI at 3T [37, 38]. To achieve the high robustness that is needed for 3D and 2D MRSI reconstruction we combine the three most powerful methods (CS, LR and SENSE) to successfully deal with the challenges of ultra-high field. We build on our previous formulation of CS-SENSE-LR reconstruction developed at 3T [14, 39], and we show that even higher spatial resolutions are reachable for brain metabolic mapping at 7T due to larger SNR and faster temporal encoding. In addition, we also extend our previous 2D CS-MRSI to 3D CS-MRSI. To the best of our knowledge, CS and LR have not been studied yet for the case of whole slab 3D 1H-MRSI in human brain either at ultra high field or at lower fields.

## II. METHODS

### A. FID-MRSI Sequence

Spectroscopic imaging was acquired with a ^1^H-FID-MRSI [9, 10] sequence implemented on a whole-body 7T MRI Magnetom scanner (Siemens, Erlangen, Germany) with a 7T-SC72CD gradient system of 70 mT/m total gradient strength and 200 mT/m/s nominal slew rate, using a 31-channel receive / birdcage transmit coil and running VB17 software. A slab selective excitation pulse of 1 ms was optimized with a Shinnar-LeRoux algorithm [40] to produce a 6.5 kHz bandwidth and was preceded by four-pulses WET [41] water suppression. The acquisition delay or echo time (TE), between the excitation and the signal acquisition was 1.3 ms in 2D and 0.9 ms in 3D. The free-induction decay (FID) was acquired with 1024 points and 8 kHz sampling rate (spectral window 26.93 ppm), which was followed by spoiler gradients (Fig. 1). The repetition time (TR) was 210 ms and the excitation flip angle was set to 15 degree to prevent *T*_1_ saturation in the metabolite signal, considering that the maximum metabolite longitudinal relaxation at 7T in the brain is 1800 ms [42]. K-space was acquired by Cartesian elliptical phase encoding, with both fully sampled and undersampling schemes.

**FIG. 1:**
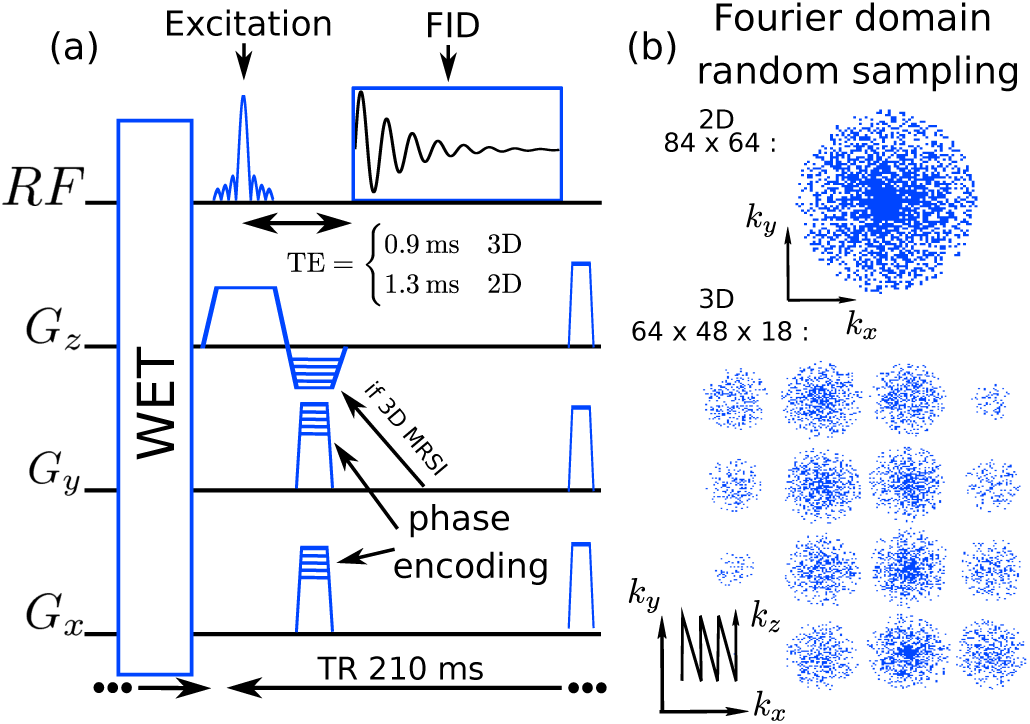
(a) Sketch of the FID-MRSI sequence for 2D and 3D. (b) Example of 2D and 3D undersampling patterns in Fourier domain.

### B. Acquisition Protocol

An anatomical 3D *T*_1_-weighted MEMPRAGE [43] volume was acquired for positioning of the 2D or 3D MRSI. The 2D MRSI was acquired with a 10 mm-thick slice and 210 *×* 160 mm field of view (FOV). Encoding matrix was 84 *×* 64 yielding a 62.5 *µ*l voxel volume (2.5 *×* 2.5 *×* 10 mm). 3D MRSI was realized with an excited slab of size 210 *×* 160 *×* 50 mm (A/P-R/L-H/F). The phase encoding FOV in the head-foot direction was set slightly larger (oversampling) to 60 mm than the 50 mm excited slab to prevent aliasing in this direction due to RF pulse excitation profile. The encoding matrix was set to 64 *×* 48 *×* 18 resulting in a 36.5 *µl* voxel volume (3.3 mm isotropic resolution). For signal referencing and to determine the coil sensitivity profiles, unsuppressed water data was acquired with the same FID-MRSI sequence and parameters but with lower resolution: 10 *×* 10 *×* 10 mm for 2D and 11.5 *×* 11.5 *×* 5 mm for 3D. The FOV size, the excited-slab thickness, the TR and flip angle were identical to main acquisition and were always acquired with a full elliptical encoding.

### C. Undersampled phase encoding

MRSI data were encoded randomly and in a sparse manner over the 3D/2D spatial Fourier domain to grant CS acceleration. Undersampling of the Cartesian encoding was performed by omitting encoding step during the sequential acquisition. The omitting pattern was computed individually in the preparation step of the running sequence. Defining the Fourier domain radius 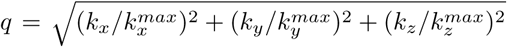, (*k*_*z*_ = 0 in 2D), the random sampling was constructed with a density distribution following *q*^−1^. The center of the Fourier space with 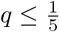 was kept fully sampled (Fig. 1).

### D. MRSI Data Processing and Reconstruction

The removal of lipid signal and residual water was performed as previously described [14]. In short, water signal remaining after the WET suppression was cleared in MRSI raw data of each coil element using the Hankel singular value decomposition (HSVD) method [44]. Afterwards, lipid suppression by metabolite-lipid orthogonality was applied to remove lipid signal in MRSI raw data for each coil element separately [14].

Following the principles of CS acceleration, MRSI data randomly undersampled must be reconstructed with a model that imposes sparsity priors while preserving fidelity with the acquired data [23]. For this study we employed a SENSE reconstruction model for arbitrary trajectory [45, 46] including total generalized variation (TGV) as regularization that imposes sparsity in 1st and 2nd order spatial derivatives. Previous implementation of CS in MRSI also used a combination of Debauchies wavelet and total variation (TV) [28, 31] or TV combined with SENSE [29]. In addition, MRSI data were assumed to be low rank and partially separable into spatial and temporal components to enhance SNR [47]. For 2D, we used the model previously published in [14, 39] and for 3D, the same model was extended to one extra spatial dimension. We describe this model shortly hereafter. MRSI raw data measured by phased-array coil element *c* = 1, …, *N*^*c*^ at time *t* and at Fourier coordinate **k** are expressed by the forward model

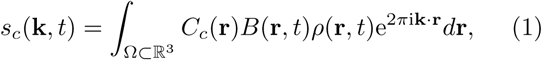

with the integration of the 3D spatial coordinates **r** over Ω, the object spatial support. The integrand is composed of the transverse magnetization *ρ*(**r**, *t*) ∈ ℂ, the coil sensitivity profiles *C*_*c*_(**r**) ∈ℂ and the spatial frequency shift 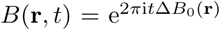 with Δ*B*_0_(**r**) the field in-homogeneity map in *Hz*. The transverse magnetization is assumed to be low-rank and separable, i.e. it can be partially separated into *K* spatial and temporal components, *U*_*n*_(**r**), *V*_*n*_(*t*) [47]:

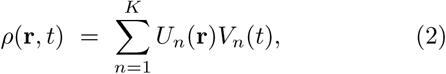

Combining (1) and (2), the forward model in vectorial notation reads

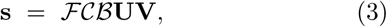

where **s** is a multi dimensional array containing the 2 or 3 spatial dimensions, the temporal dimension and one coil indexing dimension. ℱ 𝒞, and ℬ are the Fourier transform operator, the operator applying coil sensitivity profiles and the *B*_0_-inhomogeneity correction operator, respectively.

The reconstruction aims to retrieve the spatial and temporal components, *U*_*n*_(**r**), *V*_*n*_(*t*), from the sparsely sampled MRSI data *s*_*c*_(**k**, *t*). *C*_*c*_(**r**) and Δ*B*_0_(**r**) were computed from water reference acquisitions with the coils sensitivity profiles estimated using ESPIRIT [48] and *B*_0_-inhomogeneity profile estimated using *multiple signal classification algorithm* (MUSIC) [49] on the coil combined water signal spatially interpolated to the metabolite acquisition resolution.

The raw data **s** cleared of water and lipid contamination (description above) are reconstructed with a low-rank TGV model [14, 39]. The spatial and temporal components were then determined by the minimization problem including TGV spatial regularization [50]

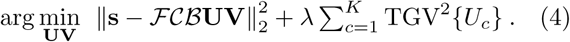

In the 2D MRSI case, a Hamming filter was integrated in the data fidelity term of the reconstruction model. The modified model eases the convergence of the reconstruction for 2D data with lower SNR while preserving effective voxel size ot the acquistion resolution. A detailed description of this filtering is given in supplementary material with some comparison on simulated, in vivo and phantom data (Supplementary and fig. S2).. The reconstruction was performed in Matlab (The MathWorks, Inc., Natick, Massachusetts, US), required a minimum of 64 RAM GB and necessitate 12hour computation time for the 3D MRSI using 8 cores of a 3.00GHz Intel(R)Xeon(R)-E5 CPU.

### E. LCModel Quantification

Metabolite concentrations were quantified using LCModel [51] and fitting the MRSI dataset *ρ* = **UV** that resulted from the optimization process in (4). A reference basis was simulated using GAMMA package [52] with acquisition parameters identical to the MRSI sequence.

The following metabolites were included in the simulated basis: N-acetylaspartate (NAA), N-acetyl aspartylglutamate (NAAG), creatine (Cr), phosphocreatine (PCr), phosphorylcholine (PCh), glycerophosphorylcholine (GPC), myo-inositol (Ins), scyllo-inositol, glutamate (Glu), glutamine, lactate, beta-glucose, alanine, taurine, aspartate, gamma-aminobutyric acid and glutathione. The removal of lipid signal by orthogonality may result in a strong baseline distortion under NAA singlet peak at 2 ppm in the form of a negative undulation. To help the LCModel quantification to cope with this distortion, a 20*Hz*-broad peak at 2 ppm and with negative phase was added to the basis. LCModel control parameters are given in the supplementary material.

The results of LCModel fitting for each voxel were further used to create spatial maps for the concentration of each metabolite. The spectral quality was assessed through the goodness of the fit and was estimated by the residuals root mean square (RMS) for each voxel.

### F. High-resolution structural and metabolic phantom

To assess precision and accuracy of the CS FID-MRSI, a high-resolution structural and metabolic phantom containing tubes of several diameters was measured with 2D FID-MRSI without CS acceleration and with a 2 mm in plane resolution. To simulate acceleration, the data were undersampled retrospectively with acceleration factor 2,3,4 and 5 following the same probability distribution in the Fourier domain as the accelerated acquisition. For comparison, a reconstruction by the forward-model adjoint operator (Fourier transform and coil combination, details in supplementary material) was also performed on fully sampled data. The geometry and molecular contrast of our custom made phantom is similar to Derenzo phantom [53, 54] used for quality control and quantification in PET molecular imaging. Our custom made phantom consists of a large cylindrical container of clear cast acrylic material (outer diameter (OD)= 15.24 cm, inner diameter (ID)= 13.33 cm, Mc-Master-Carr), 5 sets of tubes corresponding to the diameters 2, 4, 6, 8 and 10 mm, and a tube holder of size equal to the ID that is firmly fixed on the inner wall of the container. For each diameter size the set consisted of 6 tubes arranged. The tubes fixed by the holder are separated by a distance equal to twice the inner diameter of the tubes in a triangular close packed configuration as shown in fig.2. The tubes were filled with specific metabolite solutions of six different concentrations and mixtures based on their position in each size set (1-6) (top right table fig.2). Magnevist (Gd-DTPA) was added (1mL/L) in each tube to shorten T1 and create T1-weighted contrast for structural MRI. The whole tube structure was inserted in the large container which was filled with 10mM NaCl solution. The tubes and holder of the phantom were designed using 3D computer-aided design (CAD) software (rhinoceros 6.0, Robert McNeel & Associates) and fabricated using 3D printer (Formlabs, Form 2, Somerville, MA USA). The printing material was clear resin (RS-F2-GPLC-04) which is a mixture of acrylated monomers, acrylated oligomers, and photoinitiators.

**FIG. 2:**
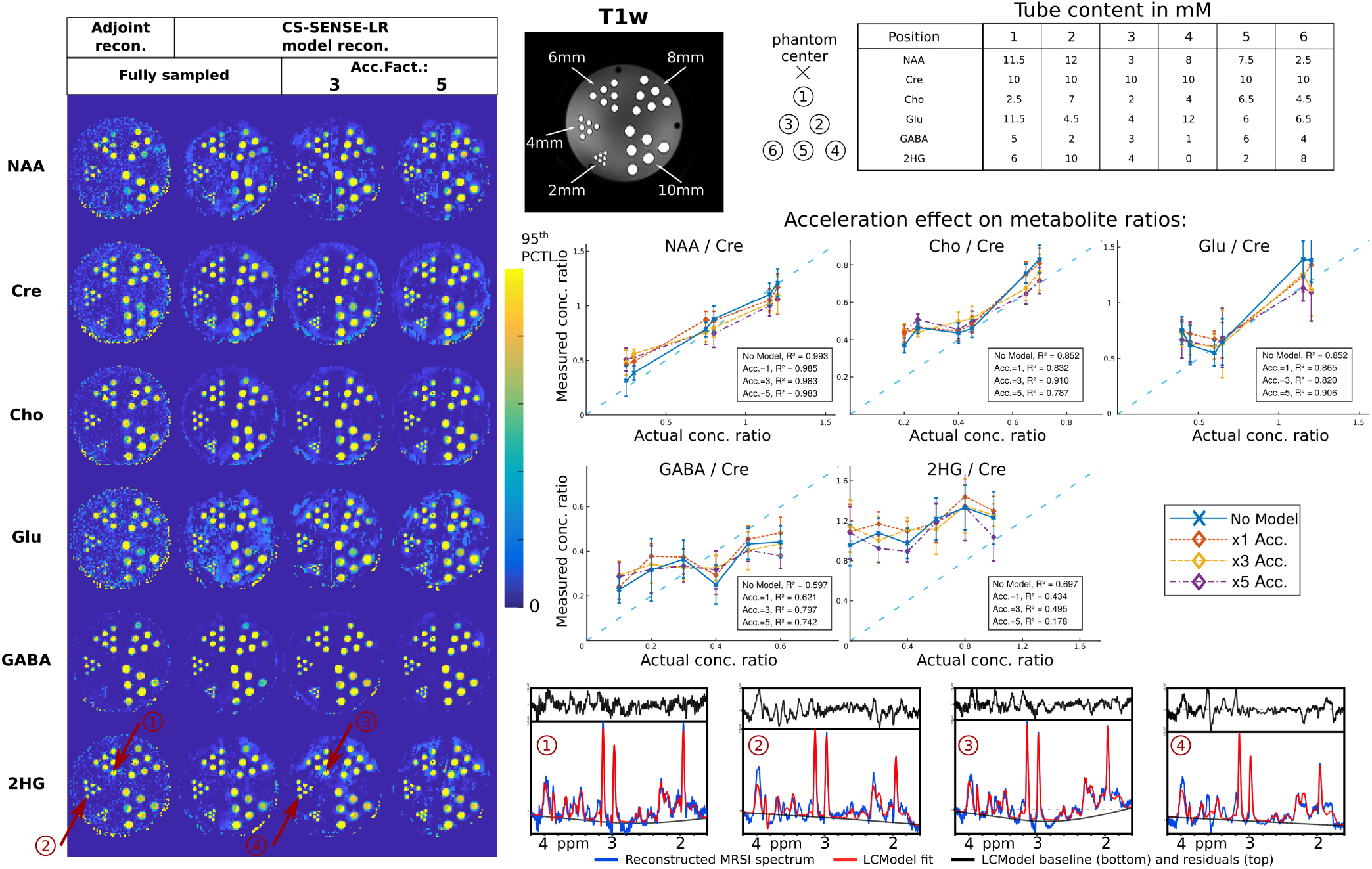
2D metabolite maps of the high structural and metabolic phantom acquired with 2 *×* 2 *×* 10 mm^3^ resolution and accelerated retrospectively with factor 1,3 or 5 (left). The actual concentration in each tube is given in the top table in mM. The tube labeling is the same for all five sizes. Right bottom, voxel-average metabolite ratio over creatine in tubes 1 to 6 of all sizes as function of the actual concentration ratio at different acceleration factors. The error bars represent the standard deviation over all the voxels within a certain tube size.

### G. Healthy Volunteer MRSI measurement

To illustrate the feasibility of the CS FID-MRSI, an acquisition protocol with 2D MRSI CS factor 2.5 (6 min) and 3D MRSI CS factor 4.5 (20 min) was acquired over 3 young healthy volunteers. In addition and to measure the effect of the CS acceleration by retrospective undersampling, a highly sampled 3D MRSI with CS 2.5 (34 min) was acquired in volunteer 1 and a fully sampled 2D MRSI (15 min) was acquired for volunteer 3. The protocol was approved by the institutional ethics committee and written informed consent was given by all subjects before participation.

## III. RESULTS

### A. high-resolution phantom

The metabolite images of the high-resolution phantom reconstructed with the CS-SENSE-R model display the tube cross-sections that are distinguishable for all diameters in comparison to the adjoint operator reconstruction that does not always permit the distiction of the 2mm tubes (fig.2). Patterns of concentrations corresponding to the metabolite content are visually observable for NAA and Cho in tubes with diameter 10, 8 and 6 mm. The Cre images and the signal variability among tubes containing all the same concentration indicate the difficulty to obtain homogeneous signal throughout the phantom for the Adjoint or the CS-SENSE-LR reconstructions. This is probably caused by the strong *B*_0_ inhomogeneities and the difficult correction of the *B*_1_ with the water signal that varies markedly in and outside the tubes due to *T*_1_ differences (see T1w water image in fig.2). As a consequence, the water signal could not be used as reference for absolute quantification but the ratio to Cre were used. In fig.2 right the measured concentration ratios versus the actual concentration ratios are displayed. The measured concentration values correspond to the voxel average in tubes from size 2 to 10 mm at one position (1-6, as label in top of fig.2) with the standard deviation as error bar. Accuracy of the results can be assessed by the distance to the diagonal (dash blue line) and the correlation coefficient square *R*^2^. NAA/Cre, Cho/Cre and Glu/Cre show good agreement between measured ratio and actual ratios for both the CS-SENSE-LR and the adjoint reconstructions although low concentration of Cho or Glu seems to be overestimated. The results for GABA/Cre and 2HG/Cre are notably less accurate with markedly low correlation coefficients and illustrate some difficulties to quantify metabolites that strongly overlap and have low signal. There is large overlap between 2HG, GABA and glutamate in short echo FID spectra which cannot be resolved by LCModel fitting. This is obvious in the case of adjoint reconstruction which shows the same fitting pattern, and hence it is a proof that it is not a result of the CS-SENSE-LR reconstruction. In order to better separate overlapping metabolites optimized RF excitation schemes that manipulate spin evolution would need to be used. But more importantly, the results are practically unchanged when using the CS-SENSE-LR model and increasing acceleration factors, which confirms the absence of reconstruction artifacts.

### B. Healthy Volunteer MRSI measurement

The 2D metabolite maps reconstructed from the 3 volunteer measurements with CS factor 2.5 are shown in fig.3 for NAA, tCr, Cho, Glu, Ins and NAAG with the *T*_1_-weighted image corresponding to the slice location. The maps reveal spatial anatomical patterns that are specific for each metabolite distribution in the brain and that are common to all three volunteer datasets. NAA distribution is homogeneous throughout the slice whereas tCr concentration is higher in grey matter (GM) than white matter (WM). Cho map exhibits a major concentration in the frontal WM while concentrations are low in the occipital lobe. The strongest GM/WM contrast is present in Glu concentration maps where the cortex can be distinguished. NAAG is clearly only detected in WM although it is the least accurate map due to NAAG’s low signal and overlap with NAA.

**FIG. 3:**
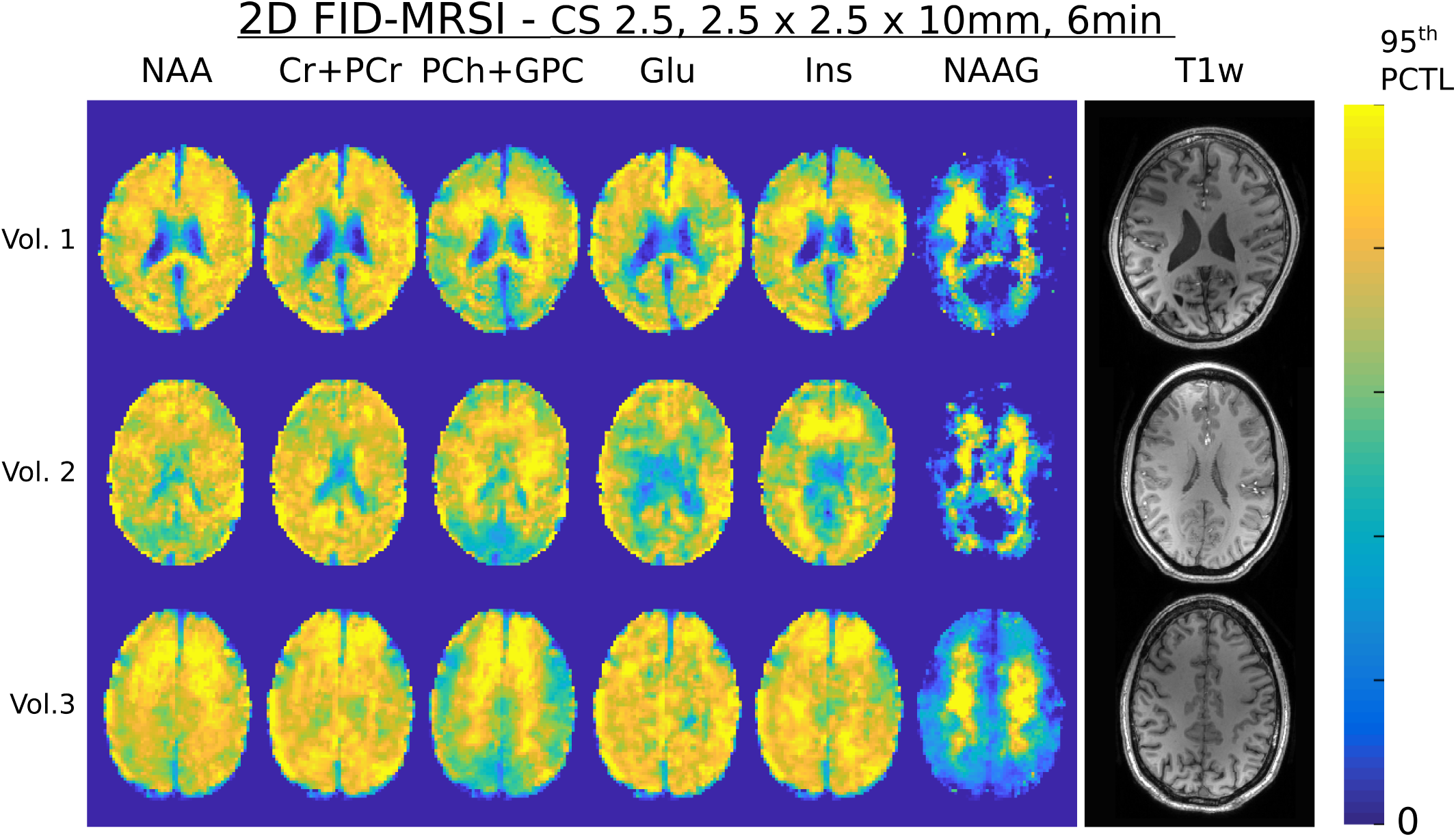
2D CS-FID-MRSI metabolite maps of three healthy volunteers measured with acceleration factor of 2.5 for a total acquisition time of 6 min. The colour scale goes from 0 till the 95th percentile for each metabolite separately. The T1-weighted anatomical image corresponding to the MRSI slice is shown to the right.

The same spatial features characterizing metabolite distributions can be observed in the 3D metabolite volumes. These were measured on the same three volunteers with 3D FID-MRSI accelerated with CS factor 4.5 and are presented in fig.4. The *T*_1_-weighted images show the anatomical content of the slab. The metabolite volumes show uniform signal and quality over the whole brain slab in spite of strong *B*_1_ and *B*_0_ inhomogeneities thanks to the intensity and frequency correction using the water-unsuppressed measurement as reference.

**FIG. 4:**
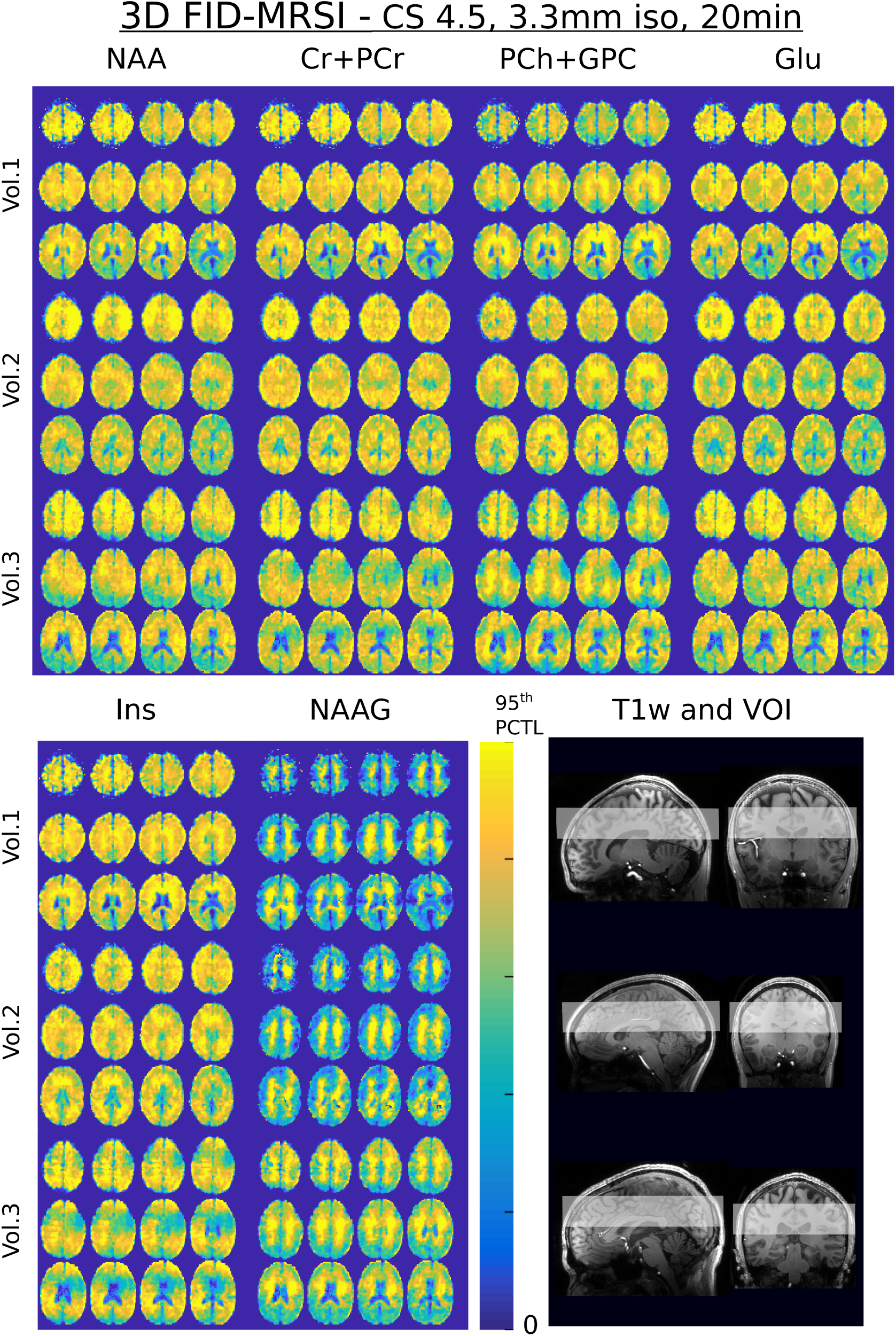
3D CS-FID-MRSI metabolite volumes of three healthy volunteers performed with acceleration factor of 4.5 for a total acquisition time of 20min. The colour scale goes from 0 till the 95th percentile for each metabolite separately. The T1-weighted anatomical images to the right show the MRSI slab positions.

### C. Retrospective Acceleration on Healthy Volunteer MRSI measurement

To evaluate the performance of CS acceleration on reconstructed metabolite maps and volumes in volunteers, 2D fully sampled or 3D highly sampled FID-MRSI data were reconstructed with several acceleration factors by retrospective undersampling (fig.5 and 6). Qualitative analysis of the 2D metabolite maps (fig.5) show almost no visible effect of the acceleration up to a factor CS=3. For higher acceleration a blurring effect is visible with a loss of small contrasts, consequence of too strong Fourier undersampling. Same effect is observable on retrospective acceleration performed on simulated data in supplementary material and figure S7. This loss of resolution was previously reported for high accelerated CS MRI [25, 46] .The sample spectra at three locations demonstrate that spectral quality is not affected by Fourier undersampling even for the acceleration factor 5. This is confirmed by the fitting residuals RMS shown in the bottom right plot that exhibit no change with the acceleration.

**FIG. 5:**
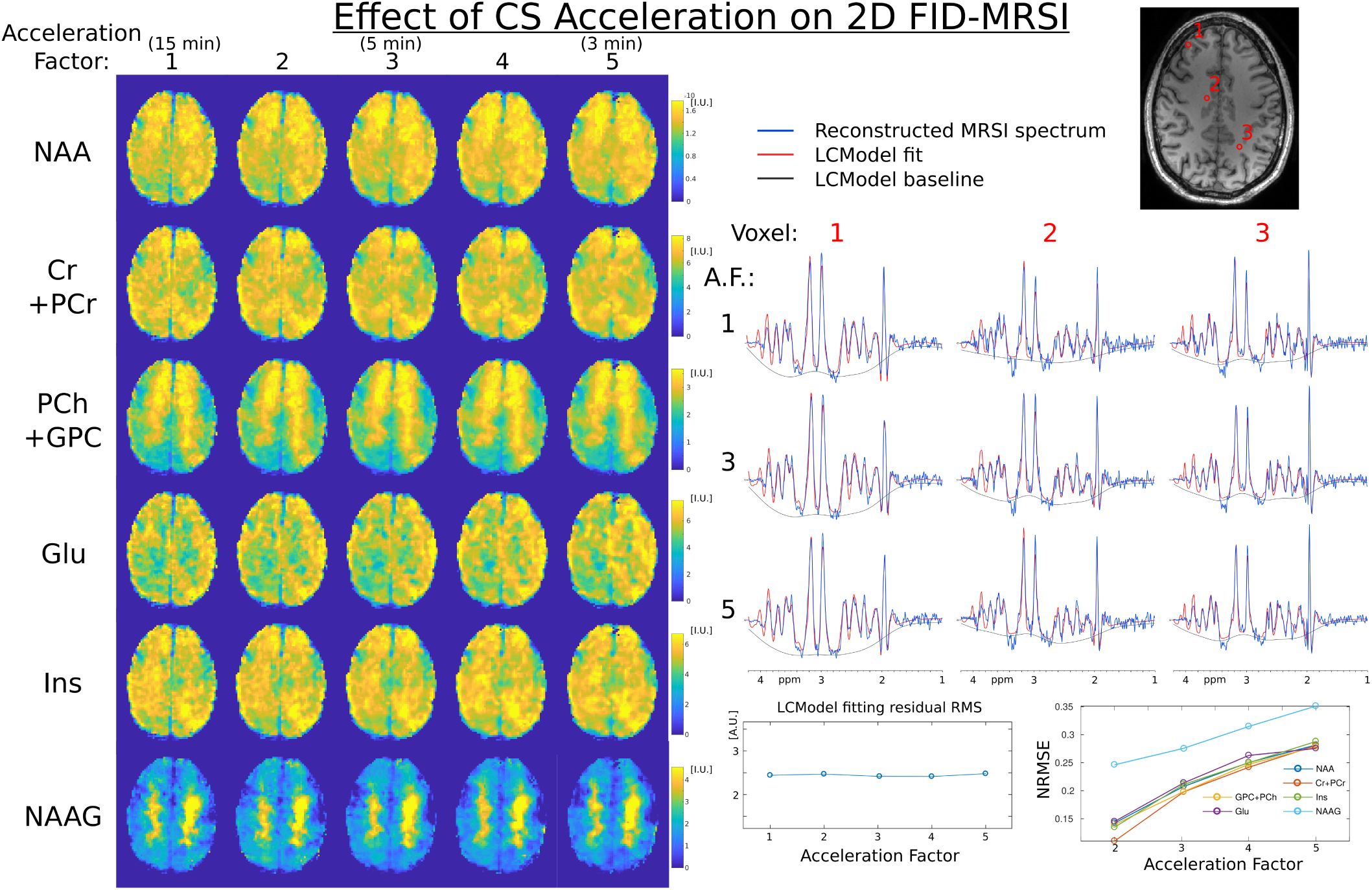
Left, 2D 2.5mm metabolite maps measured with CS-FID-MRSI and accelerated retrospectively with several acceleration factors. Scales are in institutional unit (common to all metabolites). Right, Corresponding T1-weighted image and three sample spectra are shown for acceleration factor 1,3 and 5. Bottom right, the normalized root mean square error of each metabolite relative to the fully sampled map (A.F.=1) versus the acceleration factor

The baseline distortion produced by the lipid removal by orthogonality and visible at 2 ppm is properly corrected by the inverse peak included in the lcmodel basis. Decomposed LCModel fits are shown in supplementary material fig.S6 where the baseline distortion is fitted by the ‘Inverse Broad Peak’.. This point is further considered in the discussion. A normalized root mean square error (NRMSE) over the brain for each metabolite is computed for each acceleration factor relative to the no-acceleration dataset. Let 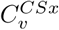 being the metabolite concentration at voxel *v* for CS acceleration factor *x*, the NRMSE <u>fo</u>r the metabolite map reads 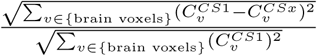. The NRMSE increases linearly or slower with the acceleration factor in agreement with previously published work [31, 55, 56] and tends to be greater for metabolite with low signal, i.e. NAAG or Glu. The effect of CS acceleration are illustrated on 3D metabolite volumes of one volunteer in fig. 6 in an orthogonal view. While acceleration of the MRSI data was progressively increased up to 6.5, only a slight loss of fine detailed contrast can be observed and no effect is visible on the three spectra shown. Quantitative error analysis with NRMSE show a linear behavior similar to the 2D case and the fitting residuals RMS stay constant when increasing the acceleration factor. The difference in SNR between 2D and 3D is particularly visible on the sample spectra.

**FIG. 6:**
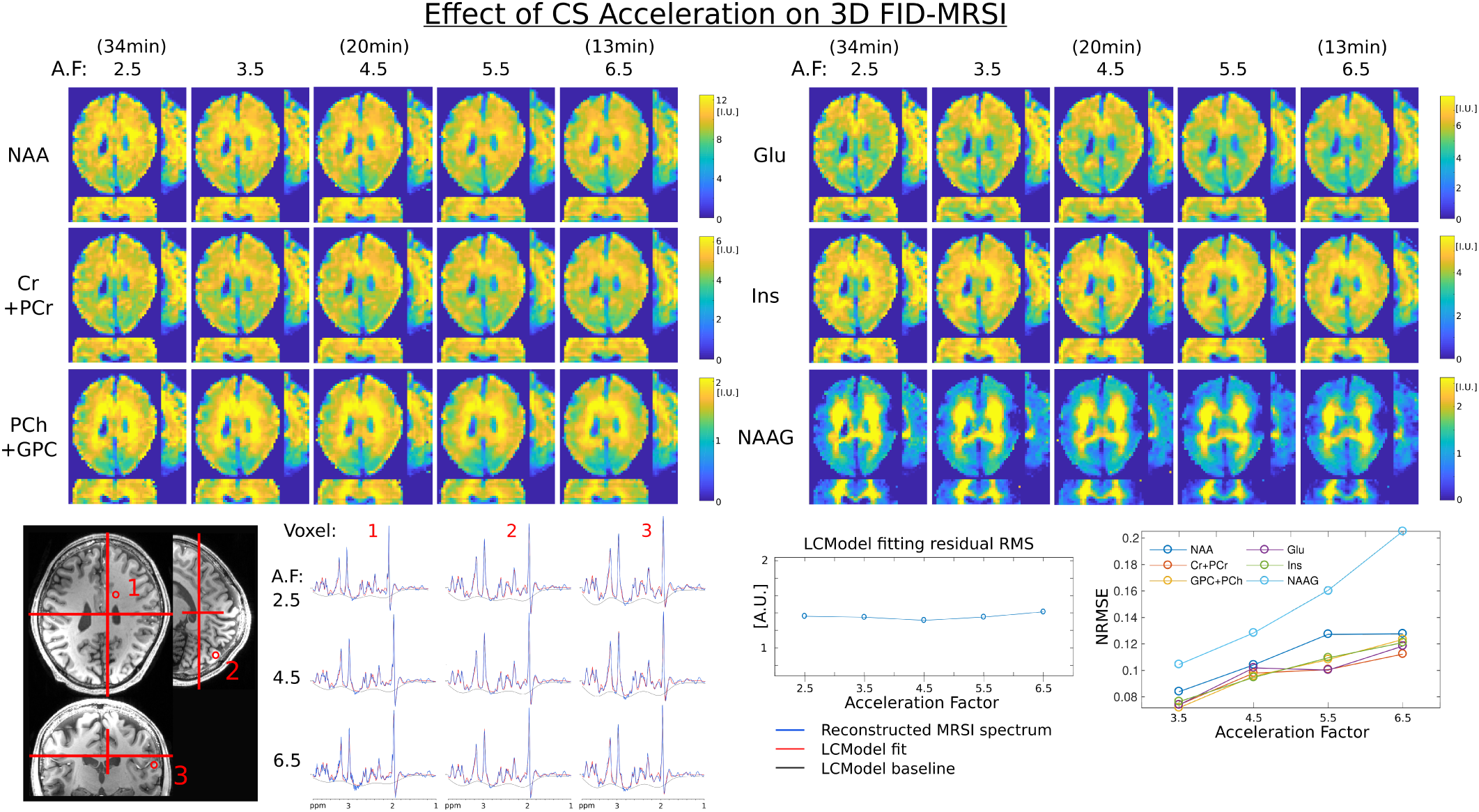
Top, 3D 3.3mm isotropic metabolite volumes acquired with CS-FID-MRSI and different acceleration factors. Bottom left, T1-weighted images of the corresponding metabolite slices in the three direction are shown next to three sample spectra at acceleration factor 2.5, 4.5 and 6.5. Bottom right, the normalized root mean square error of each metabolite relative to A.F.=2.5 versus the higher acceleration factors.

## IV. DISCUSSION

Whole-brain high-resolution 2D/3D MRSI at ultra high field can be performed with greatly reduced acquisition time and high sensitivity by combining FID-MRSI acquisition with CS-SENSE-LR acceleration and reconstruction. This achievement was enabled by the use of several technical improvements. The TR of the FID-MRSI sequence was minimized to 210 ms. Cartesian phase encoding was accelerated by random undersampling. The CS-SENSE-LR model enables accurate MRSI data reconstruction while enhancing signal with LR constraint.

The results of retrospective acceleration applied on the FID-MRSI data in 2D or 3D (fig.2, 5 and fig.6) showed that the CS-SENSE-LR model allows for accurate reconstruction albeit random undersampling is performed by factor 3 or more. For 3D MRSI (fig.6), the reference dataset was not fully sampled but was acquired with a 2.5 acceleration factor. A fully sampled 3D acquisition would be lengthy (85 min) with a high risk of head motion and large scanner frequency drift. It was assumed that considering the CS results in 2D (fig. 2, 5 and [14]), a acceleration factor of 2.5 in 3D should contain minimal acceleration distortion and represent a good starting point for retrospective acceleration.

For the highest acceleration (*≥* 4 in 2D and *≥*6 in 3D), a loss in fine details of the metabolite maps is observed. The signal containing these sharp contrasts is stored in the high spatial frequencies located in the outer Fourier space, precisely where most of the sampling points are removed. It is therefore consistent that with utmost undersampling, small contrast features are lost. This finding is consistent with the literature [25, 46] where strong CS MRI acceleration results in noticeable blurriness or loss of image resolution. These image alteration at high CS acceleration can affect small features or large structure differently and therefore cannot be described as a unique increase of the effective voxel size.

Nassirpour *et al*. observed an increasing contamination by lipid signal for higher acceleration [57] but that was not the case in our results and is probably due to the fact that in our proposed pipeline, lipid and water suppression take place before the reconstruction whereas in [57] these are performed after reconstructing the MRSI. In particular, in the previous work [57] which investigated separately the performance of low-rank and CS reconstruction, it was observed that low-rank method provided less lipid artifacts while CS provided lower CRLB for metabolite fitting. Hence, by combining LR and CS in our reconstruction we can take advantage of both properties to improve data quality and quantification. Spectral quality observable in fig.5 and 6 with the different spectra and assessed by the fitting residuals is unaffected by CS acceleration, in agreement with previous observations [57].

2D MRSI results may suffer from lower SNR in comparison to 3D MRSI even though voxel volume is larger in 2D case. This is a direct consequence of the larger volume excited by the 3D sequence and might favor a 3D over 2D multislices MRSI acquisition for particular application that requires high SNR for detection of low signal metabolite. Acceleration factor of 3 in 2D (5 min) and 4.5 in 3D (20 min) were found to be the best compromise with a strong reduction in acquisition time while presenting minimum effects on metabolite maps. The optimal acceleration factor of 3 found here for 2D and 2.5 mm in-place resolution compare to the factor of 4 in 2D and 3.1 mm resolution at 9.4T found in [57]. The difference might be explained by higher SNR at 9.4T but also by the qualitative determination of the factor. Compared to our previous work [14], we were able to increase the matrix size and reduce acquisition time in 2D MRSI from an resolution of 3.3 *×* 3.3 *×* 10*mm*^3^ in 11 min at 3T to 2.5 *×* 2.5 *×* 10*mm*^3^ in 6 min at 7T thanks to higher SNR and shorter possible TR. The accelerations obtained with CS for our 2D protocol are similar to accelerations obtained for similar protocols by parallel imaging [19, 58], with the added benefit that CS-SENSE-LR reconstruction provides effective spectral denoising and less structured artifacts from undersampling. In addition, the reconstruction presented here has the advantage to produce metabolite image with an effective voxel size identical to the nominal size for fully sampled dataset without acceleration as illustrated with the high resolution structural phantom (fig.2) and on simulated data (fig.S2). When acceleration by random undersampling is performed, a loss of resolution can be visible and compare to the gfactor noise increase for GRAPPA acceleration..

The original 2D method was also successfully extended to 3D here while benefiting from 7T advantages to reach an acquisition resolution of 3.3 mm^3^ in 20 min. Our 3D protocol using CS to acquire a matrix of 64 *×* 48 *×* 18 in 20 min compares well with a recently published 3D protocol [59] using concentric rings to acquire a matrix of 80 *×* 80 *×* 47 in 15 min or 64 *×* 64 *×* 39 in 9 min. Additional acceleration of our 3D CS-MRSI protocol could be obtained by combining CS with non-cartesian trajectories as shown at 3T for MRSI with PRESS excitation [29] or in the case of combining EPSI with LR as perfomed with SPICE [37], and similar to the accelerations obtained at 7T by non-cartesian GRAPPA [21].

The TGV regularization parameter was adjusted to *λ* = 1 *×* 10^−3^ for the 2D reconstruction (same value as for 3T study [14]) and *λ* = 3 *×* 10^−4^ for the 3D reconstruction. These same values were observed to be optimal for all 3 volunteers in 3D or 2D as shown in supplementary material (fig.S3). Therefore, no adjustment of the regularization parameter is necessary for each subject but should be slightly readjusted if acquisition parameters such as slab thickness, resolution, flip-angle or coil setup are modified. However, this is generally true not only for our reconstruction but for all model based reconstructions, including LCModel fitting (e.g. a change in sequence or *B*_0_ field requires a change in fitting basis set). The number of components in the reconstruction (*K* in (2)) was chosen as follow. As described in [14], the initial estimate of the spatial and temporal components are computed by SVD on the adjoint solution. Initial spatial and temporal components are then reviewed and *K* was qualitatively chosen as being the minimum number of components containing some signal distinguishable from noise. *K* was set to 26 for 2D and 40 for 3D.

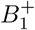 strong inhomogeneities present at ultra-high field were implicitly corrected by referencing metabolite signal with unsuppressed water data. Frequency shift yielded by *B*_0_ variation were corrected within the CS-SENSE-LR model based on a *B*_0_ fieldmap computed from the unsuppressed water data but apparent loss of signal may still occur in region where local *B*_0_ inhomogeneity is large and reduces strongly the metabolite *T*_2*_. The lipid suppression by metabolite-lipid orthogonality permits a complete removal of lipid contamination from the 2D and 3D MRSI datasets. However, a consecutive distortion of the spectra baseline under the NAA singlet at 2 ppm may be observed. The sample spectra shown in fig.5 display this distortion with a broad baseline dip resulting in a ‘W’ shape at 2 ppm. This baseline deformation was circumvent with the introduction on extra peak in the LCModel basis at 2 ppm. This peak was introduced with a negative phase opposed to NAA singlet and with a broad 20*Hz* width to prevent cross-correlation with NAA during fitting by LCModel. As results, LCModel succeeds in close fitting of the data with a smooth baseline as shown in fig.5. This effect of the lipid suppression on spectra is an important point. Although the additional peak added to the basis copes correctly with the baseline pit, we cannot fully discard an effect on NAA quantification. This point shall be addressed more in details in a future publication.. The computation time necessary for the reconstruction of 3D high-resolution MRSI data that was about 12 hours and might be a limitating factor in the application of the technique. However, we expect that with a reconstruction code fully optimized and with specific dedicated computational resources such as graphics processing units, it could be computed in significantly shorter time such in the case of MRI [36].

To the best of our knowledge, the first 3D images of whole brain metabolite distributions were published by Duijn *et al*. [60] with low-resolution Cartesian encoding at 2T MRI scanner. These results were followed by development of fast encoding EPSI/PEPSI techniques [61–64], spiral encoding [65, 66] and rosette spectroscopic imaging [67]. Group analysis of whole brain 3D MRSI provided evidence of tissue and metabolic specific differences across the brain [68, 69]. Recent publication of metabolite distributions at ultra-high field using concentricring trajectories showed an apparent GM/WM contrast in high resolution for Cho/tCr and Glx/tCr volumes [21, 59]. To highlight the sensitivity and the reproducibility of the method presented here, these metabolite features were retrieved in both 2D and 3D maps for all 3 volunteers (fig.4 and 3) with higher contrast of GM compared to WM in tCr and Glu maps. The reconstructed NAAG maps show a typical presence in central WM as previously shown in literature [70, 71]. The Cho distribution exhibits a characteristic presence in WM with high signal in frontal lobe and low level in occipital lobe.

## V. CONCLUSION

An acquisition and reconstruction combination was proposed for CS accelerated high-resolution ^1^H-FID-MRSI at ultra high-field. The reconstruction model including LR and TGV constraints enables acceleration by phase-encoding random undersampling which is demonstrated for the first time in 3D over human brain, with little effects on metabolite maps. This MRSI method grants measurement of high-resolution whole-brain slab metabolite distributions in short acquisition times: 5 min for 2D MRSI 2.5 mm resolution and of 20 min for 3D MRSI at 3.3 mm isotropic.

## VI. ACKNOWLEDGMENTS

This research was supported by the Swiss National Science Foundation grant number: IZSEZ0 188859, and the National Institutes of Health through National Cancer Institute grant 1R01CA211080.

## References

[1] G. Oz, J. R. Alger, P. B. Barker, R. Bartha, A. Bizzi, C. Boesch, P. J. Bolan, K. M. Brindle, C. Cudalbu, A. Dinçer, et al., Radiology (2014), ISSN 1527-1315.

[2] I. Tká, P. Andersen, G. Adriany, H. Merkle, K. Uurbil, and R. Gruetter, Magnetic Resonance in Medicine (2001), ISSN 07403194.

[3] D. K. Deelchand, P.-F. V. de Moortele, G. Adriany, I. Iltis, P. Andersen, J. P. Strupp, J. Thomas Vaughan, K. Uğurbil, and P.-G. Henry, Journal of Magnetic Resonance 206, 74 (2010), ISSN 10907807.

[4] B. L. van de Bank, U. E. Emir, V. O. Boer, J. J. A. van Asten, M. C. Maas, J. P. Wijnen, H. E. Kan, G. Oz, D. W. J. Klomp, and T. W. J. Scheenen, NMR in Biomedicine 28, 306 (2015), ISSN 09523480.

[5] J. W. Pan, K. M. Lo, and H. P. Hetherington, Magnetic Resonance in Medicine (2012), ISSN 07403194.

[6] A. Lutti, J. Stadler, O. Josephs, C. Windischberger, O. Speck, J. Bernarding, C. Hutton, and N. Weiskopf, PLoS ONE (2012), ISSN 19326203.

[7] P. B. Barker and D. D. Lin, Progress in Nuclear Magnetic Resonance Spectroscopy 49, 99 (2006), ISSN 00796565.

[8] A. Henning, NeuroImage 168, 181 (2018), ISSN 10538119.

[9] A. Henning, A. Fuchs, J. B. Murdoch, and P. Boesiger, NMR Biomed 22, 683 (2009).

[10] W. Bogner, S. Gruber, S. Trattnig, and M. Chmelík, NMR in Biomedicine 25, 873 (2012), ISSN 09523480.

[11] B. Bilgic, B. Gagoski, T. Kok, and E. Adalsteinsson, Magnetic Resonance in Medicine 69, 1501 (2013).

[12] C. Ma, F. Lam, C. L. Johnson, and Z. P. Liang, Magnetic Resonance in Medicine (2016), ISSN 15222594.

[13] S. Y. Tsai, Y. R. Lin, H. Y. Lin, and F. H. Lin, Magnetic Resonance in Medicine (2019), ISSN 15222594.

[14] A. Klauser, S. Courvoisier, J. Kasten, M. Kocher, M. Guerquin-Kern, D. Van De Ville, and F. Lazeyras, Magnetic Resonance in Medicine 81, 2841 (2019), ISSN 07403194.

[15] R. Otazo, B. Mueller, K. Ugurbil, L. Wald, and S. Posse, Magnetic Resonance in Medicine (2006), ISSN 07403194.

[16] L. Hingerl, W. Bogner, P. Moser, M. Považan, G. Hangel, E. Heckova, S. Gruber, S. Trattnig, and B. Strasser, Magnetic Resonance in Medicine 79, 2874 (2018), ISSN 15222594.

[17] C. Schirda, T. Zhao, H. Hetherington, V. Yushmanov, and J. Pan, in ISMRM annual meeting (2016), p. 2351.

[18] L. Valkovič, M. Chmelík, M. Meyerspeer, B. Gagoski, C. T. Rodgers, M. Krššak, O. C. Andronesi, S. Trattnig, and W. Bogner, NMR in Biomedicine 29, 1825 (2016), ISSN 09523480.

[19] B. Strasser, M. Považan, G. Hangel, L. Hingerl, M. Chmelík, S. Gruber, S. Trattnig, and W. Bogner, Magnetic Resonance in Medicine (2017), ISSN 15222594.

[20] G. Hangel, B. Strasser, M. Považan, E. Heckova, L. Hingerl, R. Boubela, S. Gruber, S. Trattnig, and W. Bogner, NeuroImage 168, 199 (2018), ISSN 1053-8119.

[21] P. Moser, W. Bogner, L. Hingerl, E. Heckova, G. Hangel, S. Motyka, S. Trattnig, and B. Strasser, Magnetic Resonance in Medicine 82, 1587 (2019), ISSN 0740-3194.

[22] D. Donoho, IEEE Transactions on Information Theory 52, 1289 (2006), ISSN 0018-9448.

[23] L. Michael, D. David, and P. J. M., Magnetic Resonance in Medicine 58, 1182 (2007).

[24] K. T. Block, M. Uecker, and J. Frahm, Magn Reson Med 57, 1086 (2007).

[25] S. D. Sharma, C. L. Fong, B. S. Tzung, M. Law, and K. S. Nayak, Investigative radiology 48, 638 (2013), ISSN 1536-0210.

[26] B. M. A. Delattre, S. Boudabbous, C. Hansen, A. Nero- ladaki, A.-L. Hachulla, and M. I. Vargas, European Radiology 30, 308 (2020), ISSN 0938-7994.

[27] R. Otazo, D. K. Sodickson, A. Yoshimoto, and S. Posse, in ISMRM annual meeting (2009), p. 331.

[28] S. Geethanath, H.-M. Baek, S. K. Ganji, Y. Ding, E. A. Maher, R. D. Sims, C. Choi, M. A. Lewis, and V. D. Kodibagkar, Radiology 262, 985 (2012), ISSN 0033-8419.

[29] I. Chatnuntawech, B. Gagoski, B. Bilgic, S. F. Cauley, K. Setsompop, and E. Adalsteinsson, Magnetic Resonance in Medicine 74, 13 (2015), ISSN 07403194.

[30] S. D. Sharma, C. L. Fong, B. S. Tzung, M. Law, and K. S. Nayak, Magnetic Resonance in Medicine 75, 42 (2016), ISSN 07403194.

[31] S. Nassirpour, P. Chang, N. Avdievitch, and A. Henning, Magnetic Resonance in Medicine (2018), ISSN 15222594.

[32] H. Y. Chen, P. E. Larson, J. W. Gordon, R. A. Bok, M. Ferrone, M. van Criekinge, L. Carvajal, P. Cao, J. M. Pauly, A. B. Kerr, et al., Magnetic Resonance in Medicine (2018), ISSN 15222594.

[33] G. H. Hatay, M. Yildirim, and E. Ozturk-Isik, Medical and Biological Engineering and Computing (2017), ISSN 17410444.

[34] G. Hangel, B. Strasser, M. Považan, S. Gruber, M. Chmelík, M. Gajdošík, S. Trattnig, and W. Bogner, NMR in Biomedicine 28, 1413 (2015).

[35] P. Balchandani and D. Spielman, Magn Reson Med 59, 980 (2008).

[36] M. Murphy, M. Alley, J. Demmel, K. Keutzer, S. Vasanawala, and M. Lustig, IEEE Transactions on Medical Imaging 31, 1250 (2012), ISSN 0278-0062.

[37] F. Lam, C. Ma, B. Clifford, C. L. Johnson, and Z.-P. Liang, Magnetic Resonance in Medicine 76, 1059 (2016).

[38] I. Bhattacharya and M. Jacob, Magnetic Resonance in Medicine 78, 1267 (2016).

[39] J. Kasten, F. Lazeyras, and D. Van De Ville, Medical Imaging, IEEE Transactions on 32, 1853 (2013).

[40] J. Pauly, P. Le Roux, D. Nishimura, and A. Macovski, IEEE Trans Med Imaging 10, 53 (1991).

[41] R. J. Ogg, P. B. Kingsley, and J. S. Taylor, Journal of Magnetic RSesonance. Series B 104, 1 (1994), ISSN 10641866.

[42] Y. Li, Journal of Molecular Imaging & Dynamics 02 (2013), ISSN 21559937.

[43] A. J. van der Kouwe, T. Benner, D. H. Salat, and B. Fischl, NeuroImage (2008), ISSN 10538119.

[44] H. Barkhuijsen, R. de Beer, and D. van Ormondt, Journal of Magnetic Resonance (1969) 73, 553 (1987), ISSN 00222364.

[45] K. P. Pruessmann, M. Weiger, P. Börnert, and P. Boesiger, Magnetic Resonance in Medicine 46, 638 (2001), ISSN 07403194.

[46] D. Liang, B. Liu, J. Wang, and L. Ying, Magnetic resonance in medicine 62, 1574 (2009), ISSN 1522-2594.

[47] H. M. Nguyen, X. Peng, M. N. Do, and Z.-P. Liang, IEEE Trans Biomed Eng 60, 78 (2013).

[48] M. Uecker, P. Lai, M. J. Murphy, P. Virtue, M. Elad, J. M. Pauly, S. S. Vasanawala, and M. Lustig, Magnetic Resonance in Medicine 71, 990 (2014), ISSN 07403194, 15334406.

[49] M. H. J. Gruber and M. H. Hayes, Technometrics 39, 335 (1997), ISSN 00401706.

[50] K. Bredies, K. Kunisch, and T. Pock, SIAM Journal on Imaging Sciences 3(3), 492 (2010), ISSN 19364954.

[51] S. W. Provencher, Magnetic Resonance in Medicine 30, 672 (1993), ISSN 0740-3194.

[52] S. Smith, T. Levante, B. Meier, and R. Ernst, Journal of Magnetic Resonance, Series A 106, 75 (1994), ISSN 10641858.

[53] S. E. Derenzo, T. F. Budinger, J. L. Cahoon, R. H. Huesman, and H. G. Jackson, IEEE Transactions on Nuclear Science 24, 544 (1977), ISSN 0018-9499.

[54] B. L. Cox, S. A. Graves, M. Farhoud, T. E. Barnhart, J. J. Jeffery, K. W. Eliceiri, and R. J. Nickles, American journal of nuclear medicine and molecular imaging 6, 199 (2016).

[55] R. Otazo, D. Kim, L. Axel, and D. K. Sodickson, Magnetic Resonance in Medicine 64, 767 (2010), ISSN 07403194.

[56] J.-X. X. Wang, M. E. Merritt, A. D. Sherry, and C. R. Malloy, Magnetic Resonance in Medicine 76, /a (2016), ISSN 15222594.

[57] S. Nassirpour, P. Chang, and A. Henning, NeuroImage 183, 336 (2018), ISSN 10538119.

[58] S. Nassirpour, P. Chang, and A. Henning, NeuroImage (2018), ISSN 10959572.

[59] L. Hingerl, B. Strasser, P. Moser, G. Hangel, S. Motyka, E. Heckova, S. Gruber, S. Trattnig, and W. Bogner, Investigative Radiology p. 1 (2019), ISSN 0020-9996.

[60] J. H. Duijn, G. B. Matson, A. A. Maudsley, and M. W. Weiner, Magnetic Resonance Imaging 10, 315 (1992), ISSN 0730725X.

[61] S. Posse, C. DeCarli, and D. Le Bihan, Radiology 192, 733 (1994), ISSN 0033-8419.

[62] A. Ebel, B. J. Soher, and A. A. Maudsley, Magnetic Resonance in Medicine 46, 1072 (2001), ISSN 0740-3194.

[63] A. Ebel and N. Schuff, Magnetic Resonance in Medicine 58, 1061 (2007), ISSN 07403194.

[64] M. Sabati, J. Zhan, V. Govind, K. L. Arheart, and A. A. Maudsley, Journal of Magnetic Resonance Imaging 39, 224 (2014), ISSN 10531807.

[65] E. Adalsteinsson, P. Irarrazabal, S. Topp, C. Meyer, A. Macovski, and D. M. Spielman, Magnetic Resonance in Medicine 39, 889 (1998), ISSN 07403194.

[66] M. Esmaeili, T. F. Bathen, B. R. Rosen, and O. C. Andronesi, Magnetic Resonance in Medicine 77, 490 (2017), ISSN 15222594.

[67] C. V. Schirda, T. Zhao, V. E. Yushmanov, Y. Lee, G. R. Ghearing, F. S. Lieberman, A. Panigrahy, H. P. Hetherington, and J. W. Pan, Magnetic Resonance in Medicine 79, 2470 (2018), ISSN 15222594.

[68] A. Maudsley, C. Domenig, V. Govind, A. Darkazanli, C. Studholme, K. Arheart, and C. Bloomer, Magnetic Resonance in Medicine 61, 548 (2009), ISSN 07403194.

[69] A. Lecocq, Y. Le Fur, A. A. Maudsley, A. Le Troter, S. Sheriff, M. Sabati, M. Donnadieu, S. Confort-Gouny, P. J. Cozzone, M. Guye, et al., Journal of Magnetic Resonance Imaging 42, 280 (2015), ISSN 10531807.

[70] M. Považan, G. Hangel, B. Strasser, S. Gruber, M. Chmelík, S. Trattnig, and W. Bogner, NeuroImage 121, 126 (2015), ISSN 10538119.

[71] P. J. W. Pouwels and J. Frahm, NMR in Biomedicine 10, 73 (1997), ISSN 0952-3480.

