## Supplementary Material for "Achieving High-Resolution Whole-Brain Slab ^1^H-MRSI with Compressed-Sensing and Low-Rank Reconstruction at 7 Tesla"

Further details about the methods and results of the main manuscript are presented in this document.

#### I. MODEL ALTERNATIVES

In fig.S1, 2D MRSI dataset from volunteer 3 and the high-resolution phantom (describe in main manuscript) are reconstructed with several alternative models. With the notation defined in the methods section of the main manuscript, we recall the forward model

$$\mathbf{s} = \mathcal{F}\mathcal{C}\mathcal{B}\mathbf{U}\mathbf{V}, \text{ with the partially separable MRSI signal as } \rho = \mathbf{U}\mathbf{V}. \tag{1}$$

The fig.S1 left shows the reconstruction result for fully elliptical sampled 2D MRSI datasets. The alternative models used for reconstruction are defined as follow

| | |
|---|---|
| *Adjoint op.* | $\rho = \mathcal{B}^*\mathcal{C}^*\mathcal{F}^*\mathbf{s}$ (with $X^*$ the adjoint of the operator $X$) |
| *SENSE* | $\arg\min_\rho \|\mathbf{s} - \mathcal{F}\mathcal{C}\mathcal{B}\rho\|_2^2$ |
| *SENSE + LR* | $\arg\min_{\mathbf{UV}} \|\mathbf{s} - \mathcal{F}\mathcal{C}\mathcal{B}\mathbf{UV}\|_2^2$ |
| *SENSE + TGV* | $\arg\min_\rho \|\mathbf{s} - \mathcal{F}\mathcal{C}\mathcal{B}\rho\|_2^2 + \lambda \sum_{t=1}^{T} \mathrm{TGV}^2 \rho(\mathbf{r}, t)$ |
| *SENSE + TGV + LR* | $\arg\min_{\mathbf{UV}} \|\mathbf{s} - \mathcal{F}\mathcal{C}\mathcal{B}\mathbf{UV}\|_2^2 + \lambda \sum_{c=1}^{K} \mathrm{TGV}^2 \{U_c\}$ |

The *Adjoint op.* reconstruction is the application of the forward model adjoint operator as reconstruction, sometimes refered as *Direct Fourier transform* reconstruction. The reconstruction *SENSE + TGV + LR* is the CS-SENSE-LR reconstruction used in the original manuscript.

The fig.S1 right presents the performance of the reconstruction alternative but for data accelerated by retrospective undersampling factor 3. The *Adjoint op.* reconstruction is not performed at this point because this direct operation is not suited for undersampled dataset (missing k-space data resulting in signal aliasing).

A spectrum from a location designated by the red arrow is shown for each reconstruction alternative. In both cases, the complete model *SENSE + TGV + LR* reconstruction, show the best denoising performance for the metabolite distributions and the sample spectra. The spectral denoising effect of the CS-SENSE-LR model is less pronounced for the phantom dataset as more SNR is present in the original data. However, in the resulting metabolite maps, the reconstruction by complete model *SENSE + TGV + LR* enables the clear distinction of the smallest 2mm tubes.

#### II. FILTERED DATA FIDELITY

In fig.S2, we demonstrated that a Hamming filter applied on the data fidelity term can improve the reconstruction result. Also we show that this improvement is different than applying a spatial hamming filter on the measured data and does not increase the effective voxel size. The MRSI data were either reconsructed by adjoint operator or by the CS-SENSE-LR model reconstruction. Defining the spatial Hamming filter operator $\mathcal{H}$, the different reconstructions read

| | |
|---|---|
| *Adjoint op.* | $\rho = \mathcal{B}^*\mathcal{C}^*\mathcal{F}^*\mathbf{s}$ |
| *No Filter Model* | $\arg\min_{\mathbf{UV}} \|\mathbf{s} - \mathcal{FCB}\mathbf{UV}\|_2^2 + \lambda\sum_{c=1}^{K} \mathrm{TGV}^2\{U_c\}$ |
| *Fidelity Filter Model* | $\arg\min_{\mathbf{UV}} \|\mathcal{H}\left(\mathbf{s} - \mathcal{FCB}\mathbf{UV}\right)\|_2^2 + \lambda\sum_{c=1}^{K} \mathrm{TGV}^2\{U_c\}$ |
| *Data Filter Model* | $\arg\min_{\mathbf{UV}} \|\mathcal{H}\mathbf{s} - \mathcal{FCB}\mathbf{UV}\|_2^2 + \lambda\sum_{c=1}^{K} \mathrm{TGV}^2\{U_c\}$ |

. In the *Data Filter Model*, the Hamming filter is applied to the measured data $\mathbf{s}$ and the iterative optimization process fits $\mathbf{UV}$ to the filtered data. Therefore, the solution of the reconstruction inherits the characteristics of the usual spatially filtered data: improved SNR but increased effective voxel size. For the *Fidelity Filter Model*, the filter is applied on both term of the data fidelity. In absence of regularization ($\lambda = 0$), the solution is identical with or without $\mathcal{H}$ and is computed by the Penrose pseudo-inverse of the forward operator : $\rho = ((\mathcal{FCB})^*\mathcal{FCB})^{-1}\mathcal{FCB}^*\mathbf{s}$. For a reconstruction with regularization ($\lambda > 0$), *Fidelity Filter Model* corresponds to a filtering of the data fidelity contribution in the convergence process. Following the filter profile, the data fidelity is fully preserved at the center of the k-space but is lowered on the outer k coordinates. The convergence process is therefore less affected by low SNR at the k-space periphery. Also, the data fidelity contribution being attenuated on the outer k-space, this is equivalent to applying a stronger TGV regularization for higher spatial frequencies.

In fig.S2 left, MRSI fully-sampled data were simulated using an analytical phantom presented in [1] with an SNR level $\infty$, 4 or 2. The 3 sample spectra on the left illustrate the respective noise levels. The reconstructed Cr+PCr or Glu+Gln metabolite map was compared to the ground truth and the normalized root mean square error (NRMSE) ($= \sqrt{\sum_{v\in\{\text{brain voxels}\}}(C_v^{g.t.} - C_v)^2}/|\mathbf{C}^{g.t.}|$) and Structural Similarity Index (SSIM) were computed for each model and SNR. The result at $SNR = \infty$ show a good agreement of the *Adjoint op.*, *No Filter Model* and *Fidelity Filter model* with the ground truth. The *Data Filter model* shows more discrepancy due to the increased effective voxel size caused by the Hamming filtering applied on the data. This is clearly visible on both metabolites. As noise increases, the *Adjoint op.* reconstruction shows the lowest performance. The *Fidelity Filter model* shows the best NRMSE and SSIM, and produces less noisy metabolite maps in comparison to the *No Filter model.* The *Data Filter model* reconstruction shows better results than the *Adjoint op.* reconstruction with small apparent noise in the maps but there is a visible increase in effective voxel size in comparison with the *Fidelity Filter* and *No Filter model* reconstructions. In fig.S2 right, the three different models are compared for 2D MRSI dataset from volunteer 3 and the high-resolution phantom (describe in main manuscript) at the proposed acceleration factor of 3. The volunteer data reconstruction show the better denoising performed by the *Fidelity Filter model* in comparison to the *No Filter model.* The difference with the *Data Filter model* is more subtle and the increase of effective voxel size is more difficult to observed than in the simulated data. At the opposite, in the phantom reconstructed metabolite maps, the difference in denoising between models is not distinct but the increased effective voxel size is clearly visible over the two smallest set of tubes for the *Data Filter model* reconstructions. *Fidelity Filter model* does not show any increase in effective voxel size in comparison to the *No Filter model.*

These results demonstrate that the *Fidelity Filter model* produces the best reconstruction in noisy dataset without increasing the effective voxel size contrarily to the *Data Filter model.* The *Fidelity Filter model* was used for the reconstruction of the 2D MRSI data for the main manuscript results.

### III. LCMODEL CONTROL PARAMETERS

LCModel [2] was performed with the controls parameters given here after. No macromolecules (MM) and lipid simulated signal were fitted with LCModel (nsimul = 0). This choice was consecutive to the observation that no MM nor lipid peaks were present in the MRSI signal after lipid suppression and reconstruction. The MM/lipid signal were supposedly removed like the scalp lipid signal by the lipid suppression by orthogonality. The non trivial control parameters were:

```
sddegz = 999.
sddegp = 7.
ppmst = 4.2
ppmend = 1
nsimul = 0
hzpppm = 297.0502
echot = 1.3
dows = T
deltat = 0.000125
degzer = 0.00
degppm = 0.00
```

[1] A. Klauser, S. Courvoisier, J. Kasten, M. Kocher, M. Guerquin-Kern, D. Van De Ville, and F. Lazeyras, Magnetic Resonance in Medicine **81**, 2841 (2019), ISSN 07403194.

[2] S. W. Provencher, Magnetic Resonance in Medicine **30**, 672 (1993), ISSN 0740-3194.

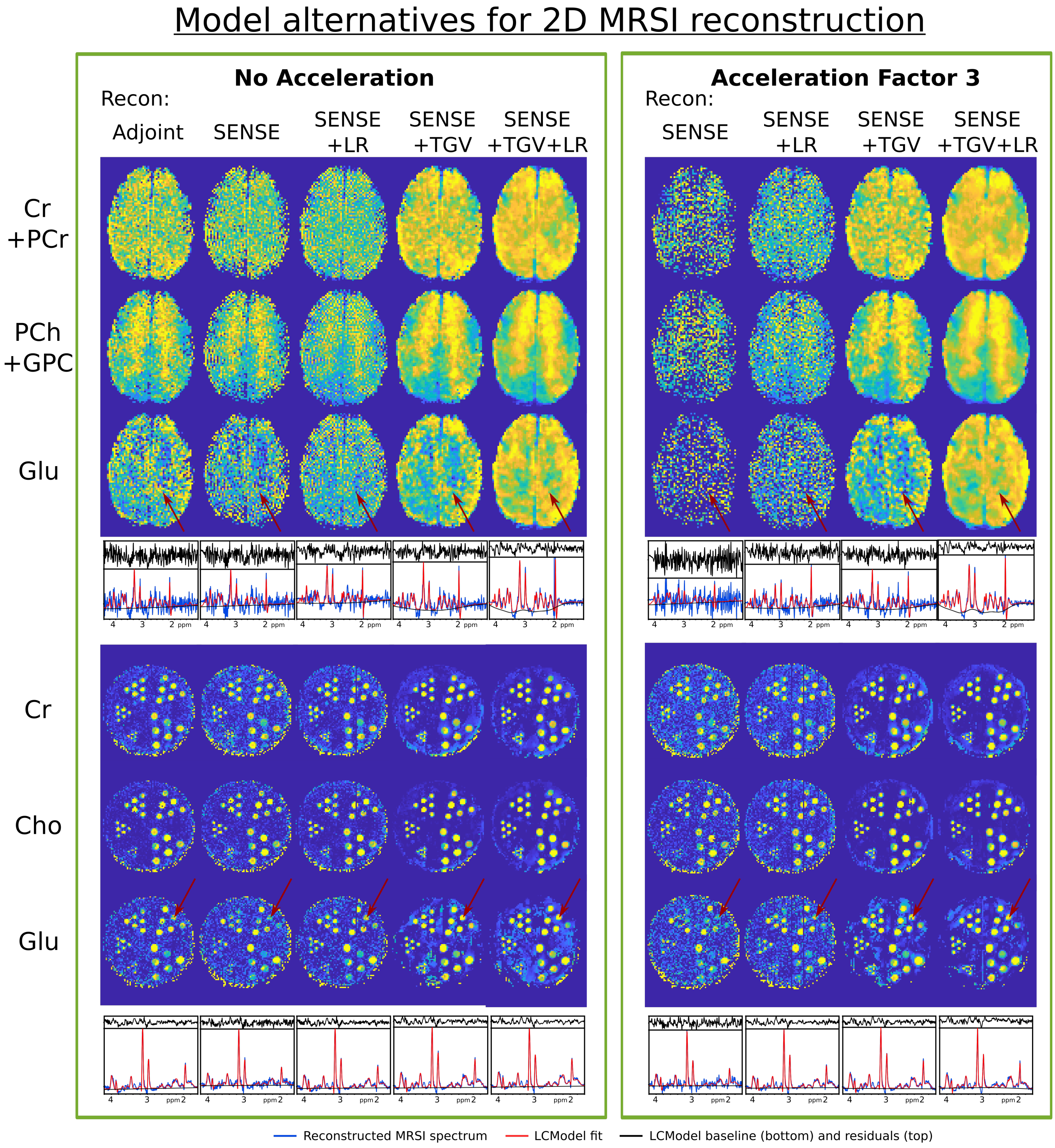


FIG. S1: Model alternatives for 2D MRSI reconstruction (see text for detailed description)

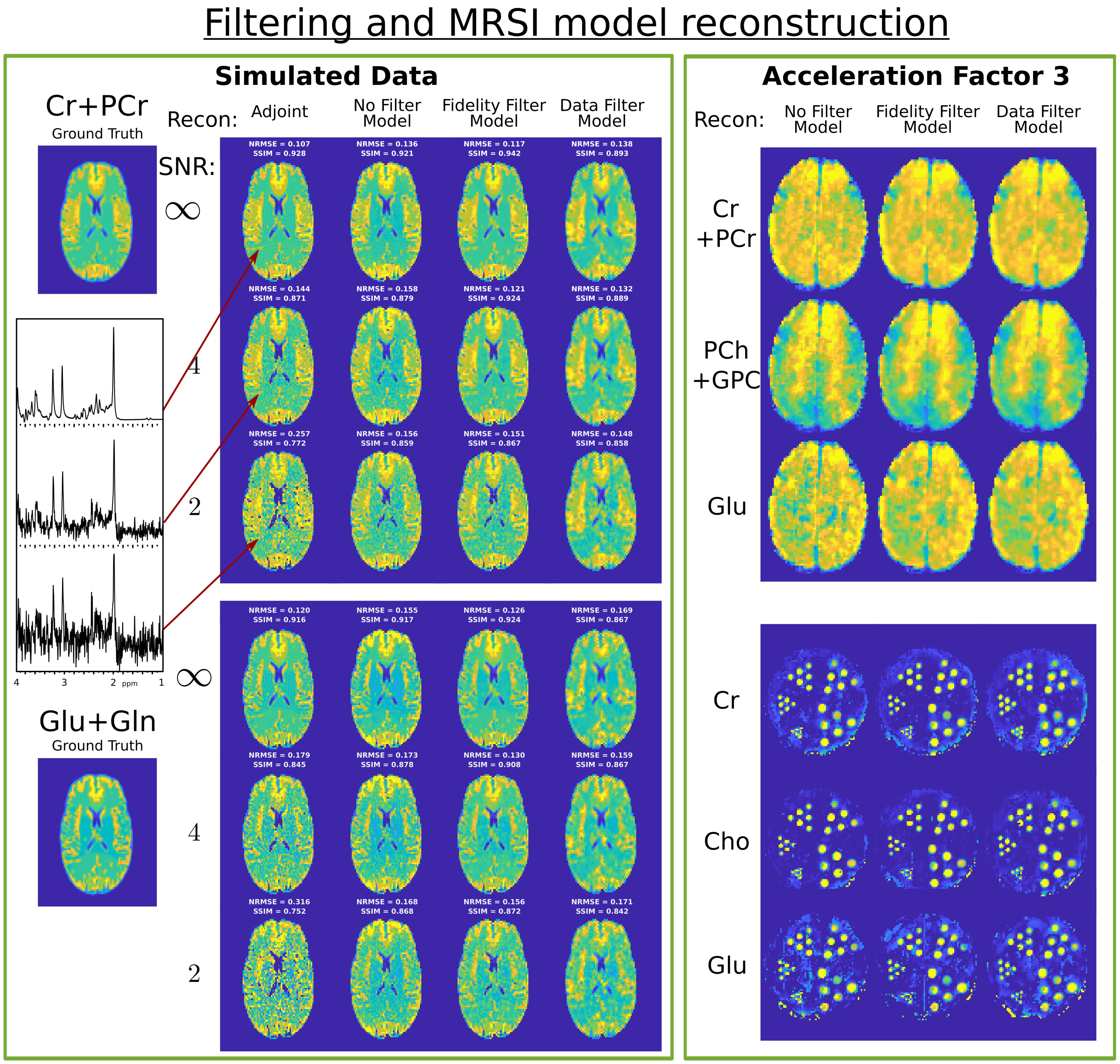


FIG. S2: Effects of different Hamming filter methods combined with the CS-SENSE-LR reconstruction model. Left, 2D MRSI were simulated with three different noise levels. The reconstructed metabolite maps of Cr+PCr and Glu+Gln were compared to the ground truth with NRMSE and SSIM. Right, in vivo and tube-phantom 2D MRSI accelerated by factor 3 were reconstructed with CS-SENSE-LR and 3 filtering methods (See text for details).

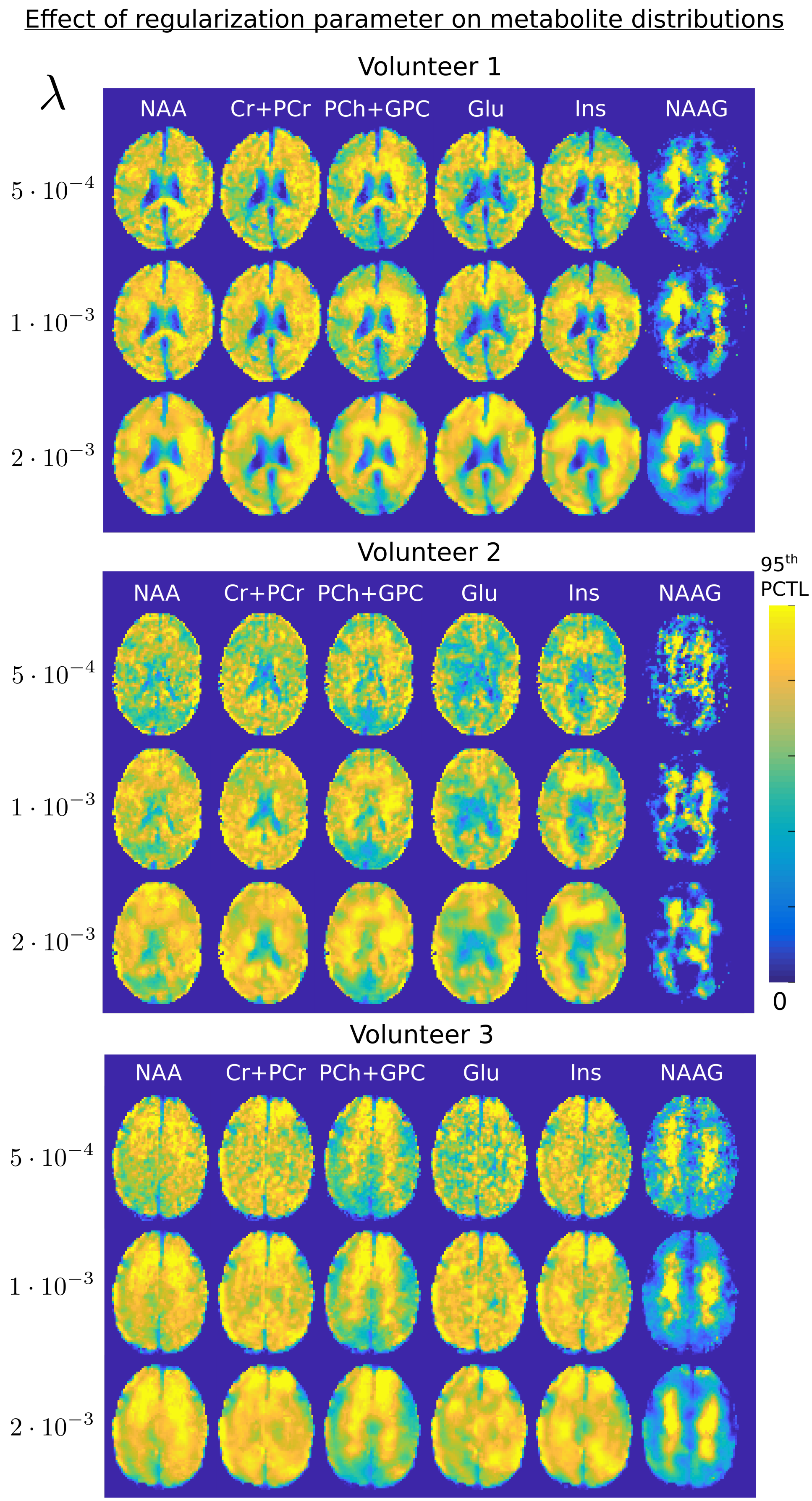


FIG. S3: Effect of the regularization parameter value on the reconstructed metabolite maps.

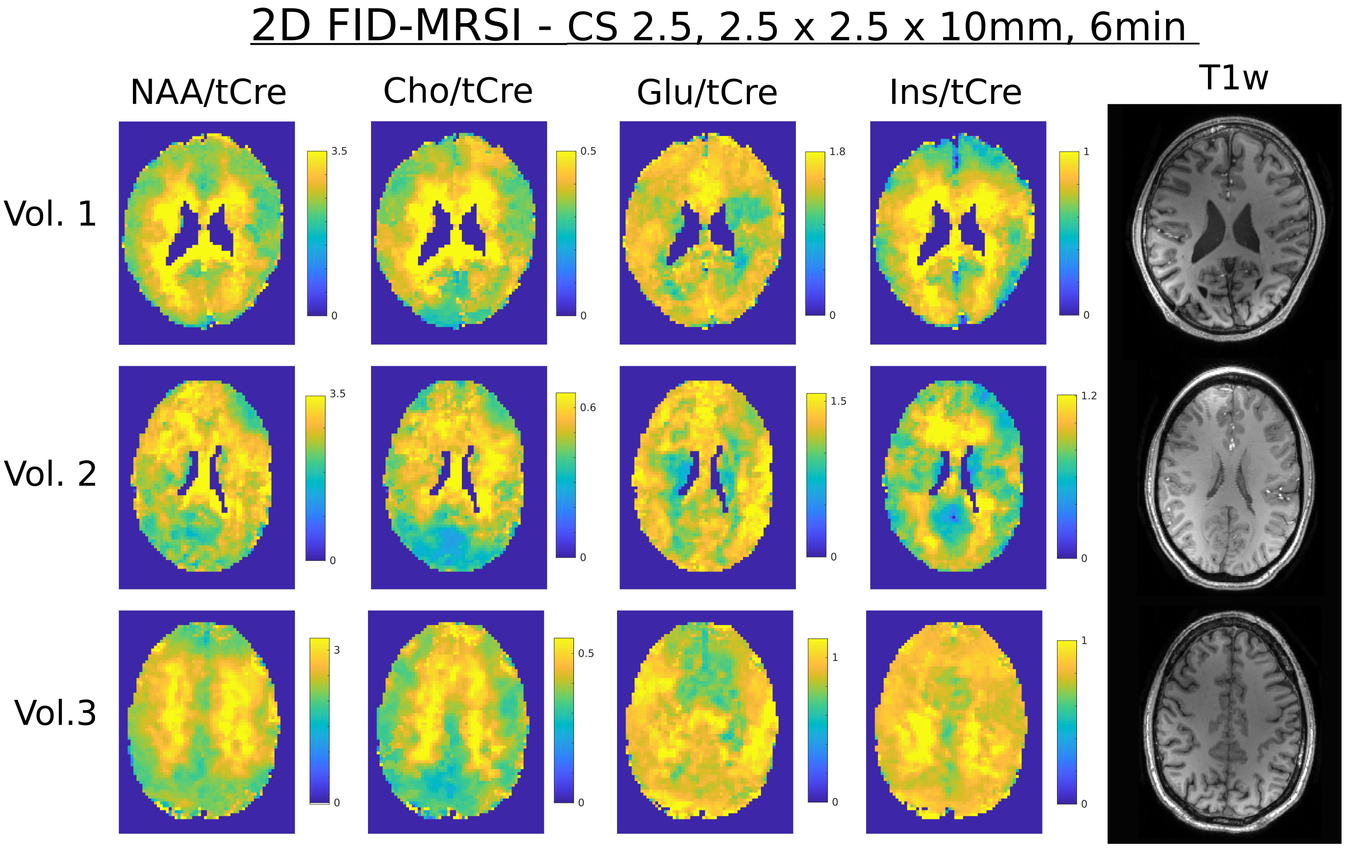


FIG. S4: 2D metabolite concentration ratio maps for the three volunteers.

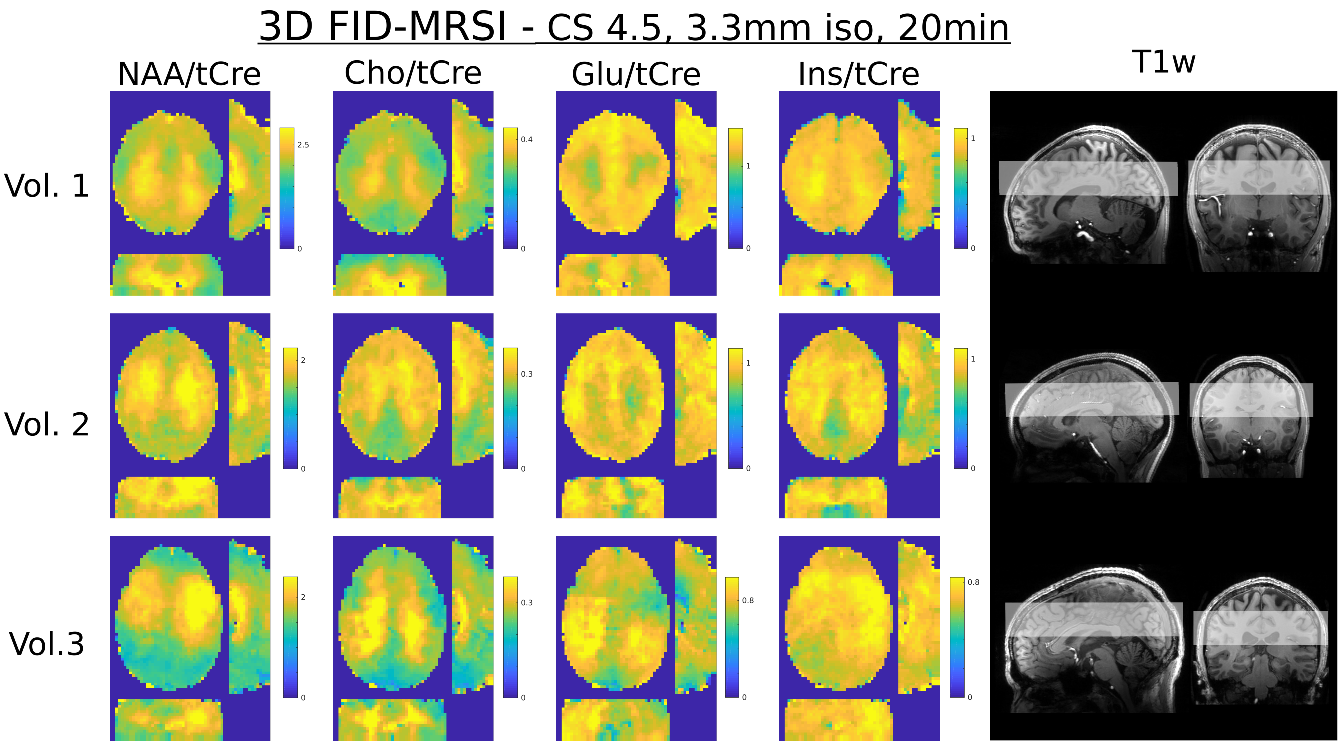


FIG. S5: 3D metabolite concentration ratio volumes for the three volunteers.

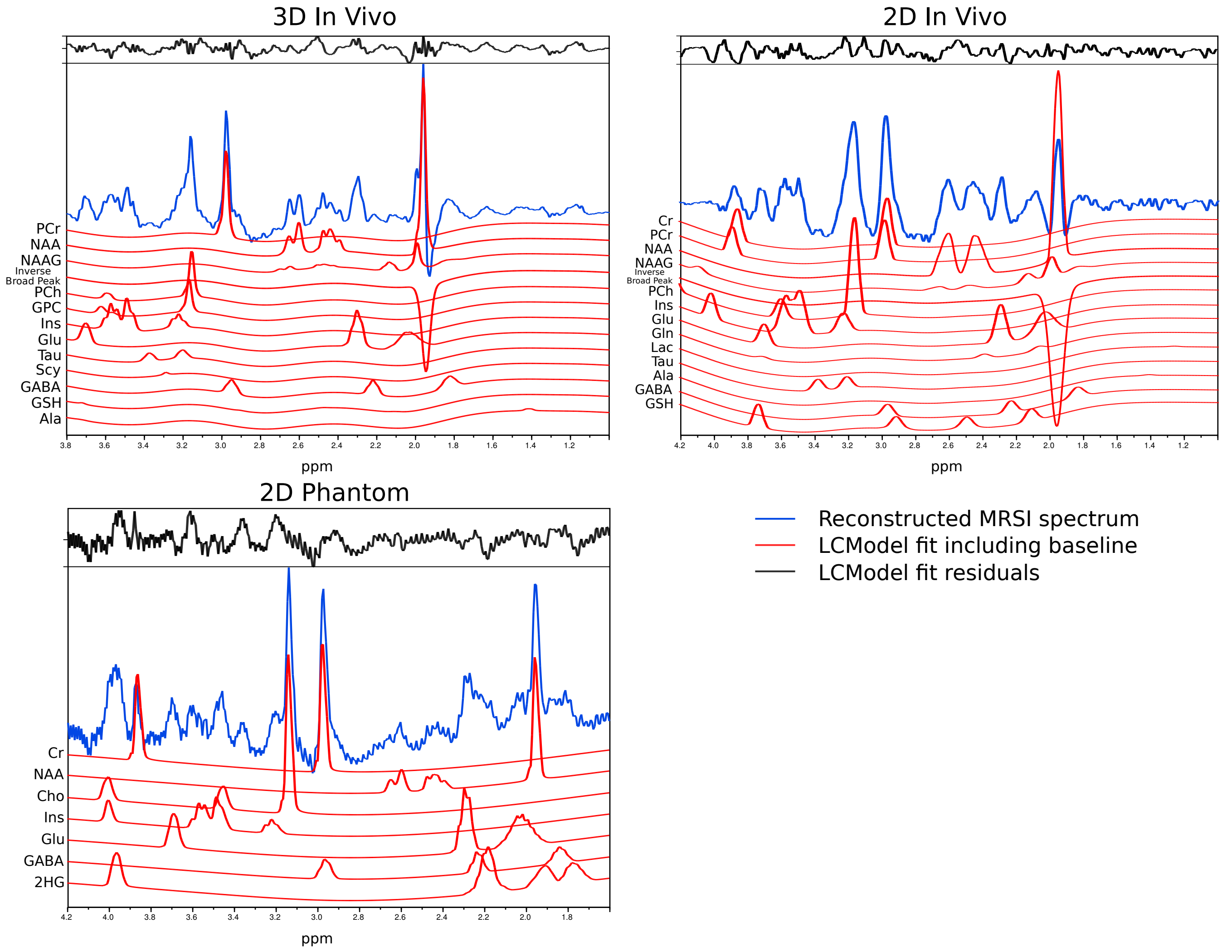


FIG. S6: Metabolite decomposition of the LCModel fitting for sample spectra. The spectra correspond to Fig.6 Voxel 1, A.F.=4.5 (3D in vivo) , Fig.5 Voxel 1, A.F.=3 (2D in vivo) and Fig.2 spectrum nb. 3 (2D phantom).

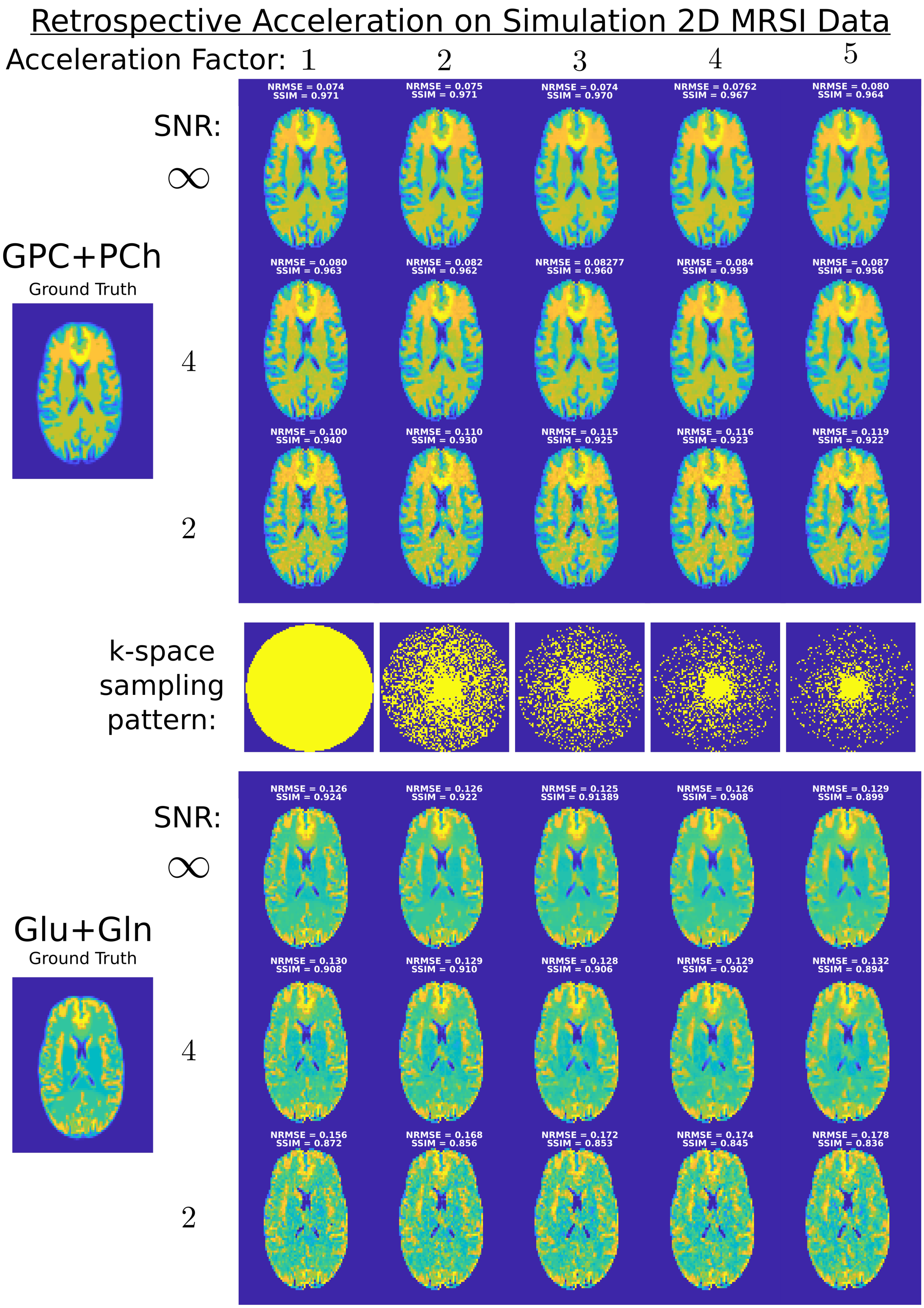


FIG. S7: Retrospective acceleration was performed on 2D MRSI simulated dataset computed with 3 levels of SNR: $\infty$, 4 or 2. An illustration of the noise level in the spectra reconstructed by adjoint op. can be found in Fig.S2. The k-space undersampling corresponding to the acceleration factor are shown in the figure center, and the GPC+PCh and Glu+Gln maps resulting from the CS-SENSE-LR reconstruction are displayed for the three levels of SNR. The ground truth concentration maps are shown on the left. For each metabolite map, the normalized root mean square error (NRMSE) and Structural Similarity Index (SSIM) were computed (see supplementary material text for details about the simulation and computation of the NRMSE and SSIM).
